# Death-associated protein kinase 3 (DAPK3) regulates the myogenic reactivity of cerebral arterioles

**DOI:** 10.1101/2022.06.10.495652

**Authors:** Sara R. Turner, David A. Carlson, Mona Chappellaz, Cindy Sutherland, Timothy A.J. Haystead, William C. Cole, Justin A. MacDonald

## Abstract

The vascular smooth muscle (VSM) of resistance blood vessels displays intrinsic autoregulatory responses to increased intraluminal pressure, the myogenic response. In the brain, the myogenic responses of cerebral arterioles are critical to homeostatic blood flow regulation. Here we provide the first evidence to link the death-associated protein kinase 3 (DAPK3) to the myogenic response of rat and human cerebral arterioles. DAPK3 is a Ser/Thr kinase involved in Ca2+− sensitization mechanisms of VSM contraction. *Ex vivo* administration of a specific DAPK3 inhibitor (i.e., HS38) could attenuate vessel constrictions invoked by serotonin as well as intraluminal pressure elevation. The HS38-dependent dilation was not associated with any change in myosin light chain (LC20) phosphorylation. The results suggest that DAPK3 does not regulate Ca2+ sensitization pathways during the myogenic response of cerebral vessels but rather operates to control the actin cytoskeleton. Finally, a slow return of myogenic tone was observed during the sustained exposure of cerebral arterioles to a suite of DAPK3 inhibitors. Recovery of tone was associated with greater LC20 phosphorylation that suggests intrinsic signaling compensation in response to attenuation of DAPK3 activity. The translational importance of DAPK3 to the human cerebral vasculature was noted, with robust expression of the protein kinase and significant HS38-dependent attenuation of myogenic reactivity found for human pial vessels.

## INTRODUCTION

The primary function of the arterioles is to regulate flow into downstream capillaries, and they achieve this through contractile tone development (Pugsley and Tabrizchi, 2000). A key component of tone development in the resistance vasculature is the myogenic response, referring to constrictions originating within the vascular smooth muscle (VSM) as a response to stretch. The vascular myogenic response is an inherent property of the resistance arteries and has been well studied in isolated pressurized vessels over the past 25 years (Hong et al., 2020). This vascular contractile response to increased luminal pressure involves membrane depolarization and Ca2+ influx, as well as several Ca^2+^ sensitization mechanisms (as reviewed in (Davis and Hill, 1999; Hill et al., 2001; Schubert et al., 2008; Cole and Welsh, 2011; El-Yazbi and Abd-Elrahman, 2017; Hong et al., 2020; Jackson, 2021)).

Despite their relatively large size, the middle cerebral arteries (MCA) and posterior cerebral arteries (PCA) display strong myogenic responses and are the primary regulators of blood flow to the cerebrum (Izzard and Heagerty, 2014). These vessels provide critical homeostatic mechanisms for the maintenance of constant blood flow to the brain, which can easily suffer damage from brief periods of either sub-or supra-optimal perfusion. The myogenic response also maintains capillary hydrostatic pressure despite fluctuations in systemic pressure and shields delicate vessels from damage caused by sudden increases in flow (Shekhar et al., 2017).

The death-associated protein kinase 3 (DAPK3, also known as zipper-interacting protein kinase or ZIPK) is a recent addition to a cadre of Ca^2+^/calmodulin-independent protein kinases involved in Ca^2+^ sensitization mechanisms of VSM (Haystead, 2005; Ihara and MacDonald, 2007). DAPK3 can phosphorylate the 20-kDa regulatory myosin light chain (LC20) (Niiro and Ikebe, 2001; Borman et al., 2002; Moffat et al., 2011; Komatsu and Ikebe, 2014), as well as the myosin phosphatase targeting subunit (MYPT1) (MacDonald et al., 2001a; Endo et al., 2004) and the 17-kDa C-kinase-potentiated inhibitor (CPI-17) of myosin phosphatase (MacDonald et al., 2001b; Xu et al., 2010). While strong evidence supports the ability of DAPK3 to regulate the contractile force of VSM (MacDonald et al., 2001a; Niiro and Ikebe, 2001; Borman et al., 2002; Endo et al., 2004; Usui et al., 2012; Carlson et al., 2013; MacDonald et al., 2016; Carlson et al., 2018), to date, there have been no examinations of DAPK3 in the vascular myogenic response. Thus, although DAPK3 can regulate VSM contractile force development under various experimental conditions, whether this signaling molecule participates in the physiological myogenic responses of cerebral arterioles to increasing intraluminal pressure remains to be investigated.

## METHODS

### Reagents

5-hydroxytryptamine (5-HT), acetylcholine (ACh) and phenylephrine (PE) were purchased from Millipore-Sigma (Oakville, ON, Canada). Anti-DAPK3 (rabbit polyclonal; ab115695) and anti-αSM-actin (rabbit polyclonal; ab5694) were obtained from Abcam (Cambridge, UK). Anti-LC20 (mouse monoclonal; sc-2839) was purchased from Santa Cruz (Dallas, TX). The HS38, HS56 and HS94 compounds were synthesized as described previously (Carlson et al., 2013; Carlson et al., 2018). Phos-Tag-acrylamide was purchased from NARD Chemicals, Inc. (Kobe City, Japan). All other chemicals were reagent grade and were obtained from Millipore-Sigma or VWR (Mississauga, ON, Canada).

### Animals

Sprague-Dawley rats (~300 g) were purchased from Charles River Laboratories (Montreal, QC) and were housed in the Animal Resource Center at the University of Calgary. Animals were housed and handled according to the standards of the Canadian Council on Animal Care and protocols reviewed by the Animal Care Committee of the Cumming School of Medicine, University of Calgary. Animals were euthanized by isoflurane inhalation to the point of being fully anesthetized, followed by decapitation by guillotine. Immediately following sacrifice, organs and tissues required for study were removed by gross dissection and placed into room temperature normal Krebs’s buffer (NKB) containing 120 mM NaCl, 25 mM NaHCO3, 4.8 mM KCl, 1.2 mM NaH2PO4, 1.2 mM MgSO4, 11 mM glucose and 1.8 mM CaCl2 (pH 7.4).

### Vessel isolation and mounting for pressure myography

PCA vessels were collected from their origin within the circle of Willis to their first major branch point, cleaned of surrounding tissue, and cut into 3 – 5 mm segments. For myography, vessels were mounted onto one glass cannula in an arteriograph chamber (Living Systems CH-1-QT; St Albans, VT, USA) and tied in place with two pieces of silk suture. To denude the vessel of endothelial cells, each vessel was passed fully onto the glass cannula, and then a stream of air bubbles was passed through the lumen of the vessel. After denuding, the other end of the vessel was mounted on the opposite cannula and tied in place with silk suture. The vessel chamber was then connected to the pressure myograph for measurement of vessel diameters (outer and inner diameters, OD and ID, respectively) via an automated edge detection system (IonOptix; Westwood, MA, USA). Vessels were briefly pressurized to 80 mmHg (~ 1 minute) to check for leaks in the system; leaky vessels were either re-tied or discarded. Pressure was then set to 10 mmHg and vessels were warmed to 37 °C by a perfusing bath of warm, aerated (with 95% air/5% CO2) NKB for a 20 minute equilibration period.

### Pressure-induced constriction protocol

Mounted PCA vessels were first tested for the presence of myogenic reactivity since the myogenic response may be lost due to mishandling during dissection and mounting. Vessels were pressurized to 80 mmHg, and the pressure was dropped back to 10 mmHg following development of myogenic constrictions and stabilization of vessel diameters. Confirmation of endothelial removal was evaluated via loss of vasodilatory response to acetylcholine (ACh, 1 μM) in vessels displaying myogenic constriction. The single step to 80 mmHg was repeated up to two additional times to confirm the stability of the myogenic response. The vessel was then subjected to pressure steps in NKB from 10 to 120 mmHg; individual pressure steps lasted for approximately 8 minutes and allowed for development of stable diameters. Pressure was then returned to 10 mmHg. In cases where DAPK3 inhibitors were tested for effects on the myogenic response, compounds were added to the bath superfusate; the vessel was then incubated at 10 mmHg for 30 minutes prior to repeating the 10 – 120 mmHg pressure steps. Pressure was then returned to 10 mmHg and the NKB, with or without inhibitor, was removed and replaced with Ca2+-free Krebs (0 Ca2+; possessing the same chemical constituents as NKB, but with no CaCl2 and addition of 2 mM ethylene glycol-bis(β-aminoethyl ether)-N,N,N’,N’-tetraacetic acid, EGTA). Data represent the average vessel diameters during stable development at each pressure step and were expressed as a percentage of the maximal passive vessel diameter in 0 Ca2+ solution to account for variations in the sizes of vessels among animals. Some vessels were flash-frozen with a dry ice – methanol bath; these were transferred to 1.5 mL microtubes, lyophilized for 16 hours, and then stored at −80 °C.

### Human cerebral vessel pressure myography and western immunoblotting

Human brain tissue samples were obtained from surgical patients following written informed consent, and in accordance with the guidelines of the Declaration of Helsinki. Ethical approval was provided by the University of Calgary and Calgary Health Region Conjoint Health Research Ethics Board (Ethics CHREB ID: 024735). Brain tissues were collected by the surgeon, placed in cold NB and then rapidly transferred to the laboratory post-operatively. Small superficial cerebral pial arteries, approximately 150 to 250 μm in diameter, were carefully dissected from surrounding tissue and cut into segments to be mounted for pressure myography. Larger superficial cerebral arteries, approximately 300 to 500 μm in diameter and 1 cm in length, were removed and flash frozen for biochemical assessments.

### Western Immunoblotting

SDS-PAGE Sample Buffer (50 μL) containing 60 mM Tris-HCl (pH 6.8), 4% (w/v) sodium dodecyl sulfate (SDS), 10 mM dithiothreitol, 10% (v/v) glycerol and 0.01% (w/v) bromophenol blue was added to a microtube containing a single PCA vessel. The tubes were briefly centrifuged (1 min; 13,000 rpm) to ensure tissues were fully submerged in Sample Buffer and were then vortexed for 16 hours at 4 °C. The samples were centrifuged again briefly (1 minute; 13,000 rpm), heated to 95 °C for 10 minutes and then stored at – 20 °C until use. Two-and three-step western immunoblotting procedures were conducted as previously described (Johnson et al., 2009). Proteins were separated on 12% SDS-PAGE gels using standard conditions. Proteins were then transferred to nitrocellulose membranes (0.2 μm pore size; Bio-Rad) at 25 V (constant voltage) for 16 hours at 4 °C using standard Tris-glycine buffers (25 mM Tris, 192 mM glycine, pH 8.3 and 20% (v/v) methanol). After blocking with 5% non-fat milk protein, membranes were sequentially incubated with primary antibody (a 1:1000 dilution) for 16 hours at 4 °C and then biotin-conjugated goat anti-rabbit IgG (Millipore-Sigma/Chemicon, Oakville, ON, Canada) secondary antibody (1:10,000 dilution) for 1 hour at room temperature. After extensive washing, membranes were incubated for 30 minutes at room temperature in HRP-conjugated streptavidin (ThermoFisher Scientific/Pierce, Waltham, MA, USA) at 1:20,000 dilution. A final washing step was completed and then membranes were developed with enhanced chemiluminescence (ECL) reagent and imaged using an LAS4000 Gel Imager with ImageQuant densitometry software (GE Healthcare, Mississauga, ON, Canada).

### LC20 Phosphorylation

Phos-tag SDS-PAGE was used to monitor LC20 phosphorylation status as previously described (MacDonald et al., 2016; Chappellaz et al., 2018). PCA tissue extracts were resolved with Phos-tag SDS-PAGE gels containing 12.5% (w/v) acrylamide, 0.42% (w/v) N,N’-methylene bisacrylamide solution with 0.1% (w/v) SDS, 60 μM MnCl2 and 30 μM Phos-tag reagent. Gels were developed at 30 mA/gel in standard Tris-glycine buffer for 70 minutes. Proteins were transferred to 0.2 μm polyvinylidene difluoride (PVDF) membranes and fixed with 0.5% (v/v) glutaraldehyde solution. Membranes were probed with primary anti-LC20 antibody (1:1000 dilution), incubated with HRP-conjugated secondary antibody (1:10,000 dilution), developed with ECL reagent and then visualized with an LAS4000 Gel Imager.

### Data Analysis

Values are presented as the mean ± S.E.M., with n indicating the number of animals (tissue experiments). Data were analyzed using Graphpad-Prism software. Multiple group comparisons were made by two-way analysis of variance (ANOVA) with Sidak’s *post hoc* test. Differences were considered statistically significant when P < 0.05.

## RESULTS

The PCA was selected as the vessel of choice to evaluate the contribution of DAPK3 to the vascular myogenic response. These arterioles, along with the MCAs, display a high degree of myogenic reactivity (Colinas et al., 2015; Abd-Elrahman et al., 2017). A concentration-response experiment was conducted to confirm the amount of HS38 required to inhibit DAPK3 in isolated pressurized vessels *ex vivo.* As shown in **Figure 1A**, vessels were pressurized to 10 mmHg and preconstricted with 1 μM 5-hydroxytryptamine (5-HT). After a stable vessel diameter was established, HS38 was added to the bath at increasing concentrations (0.1 to 20 μM; DMSO was used as a vehicle control). Vessels constricted by 17.5 ± 6.2% in response to 5-HT (relative to their initial diameter), with an average diameter change of 52.1 ± 8.8 μm (**Figure 1B**). In the vehicle controls, the vessel constriction elicited with 5-HT diminished over time; the vessels dilated by 20.8 μm or 41.4 ± 4.5% by the conclusion of the experiment. However, HS38 was found to significantly dilate the 5-HT-constricted vessel beyond that observed for the DMSO control. In this case, the maximal dilation with HS38 treatment was 45.6 μm or 87.6 ± 7.9% with a calculated EC50 of 0.65 μM.

**Figure 1.**
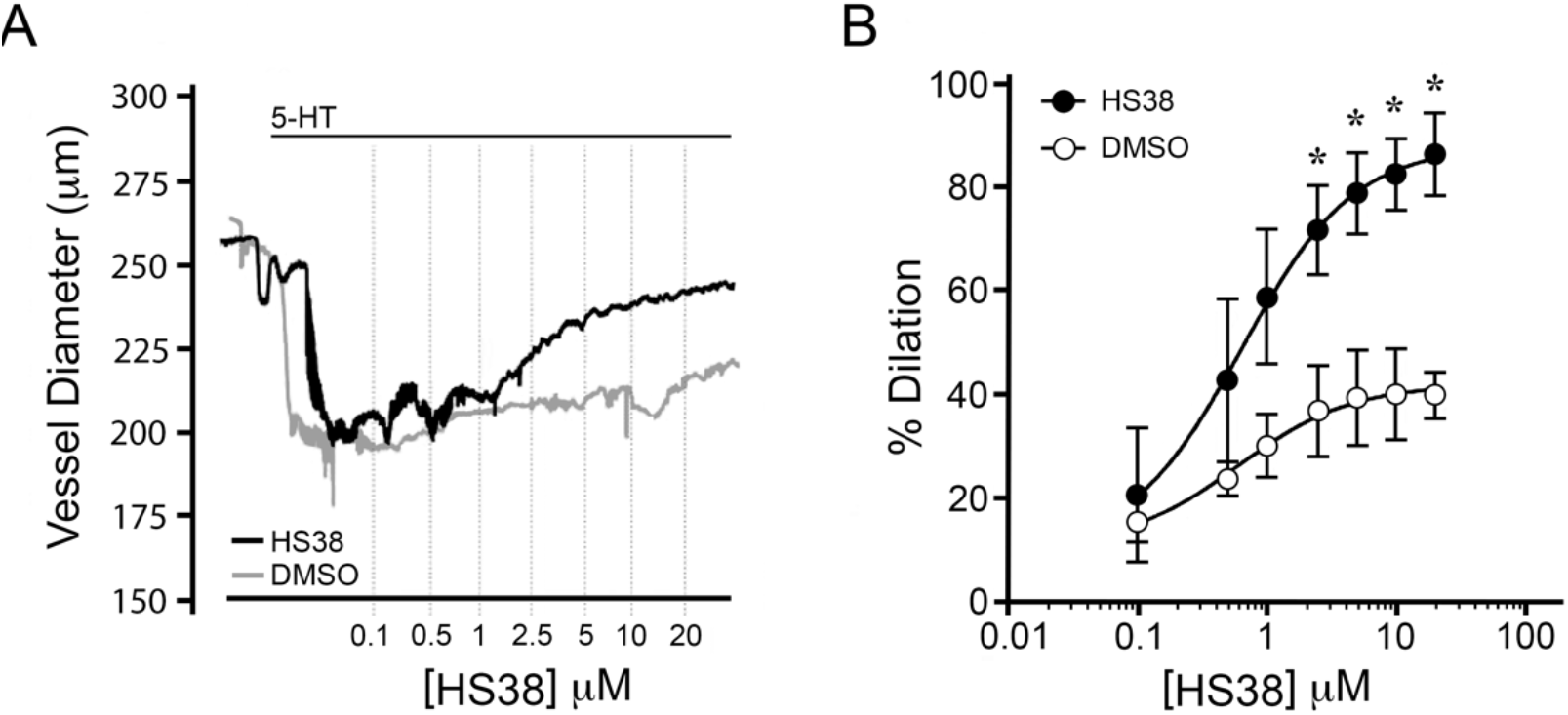
HS38 treatment relaxes rat posterior cerebral arteries constricted with serotonin. Posterior cerebral arteries were mounted at 10 mmHg and constricted with continuous exposure to serotonin hydrochloride (5-HT, 1 μM). Increasing concentrations of HS38 (0.1 – 20 μM) were applied to the bath chamber. **(A)**, representative recordings of the change in outer diameter for 5-HT constricted vessels subjected to sequential application of HS38. **(B)**, cumulative data showing vessel constriction in the presence or absence of HS38 relative to the maximum passive diameter observed in 0 Ca2+ solution. *-significantly different from vessel constriction observed with vehicle treatment (two-way ANOVA with Sidak’s *post hoc* test, p < 0.05).

Knowing that the application of HS38 was able to attenuate the constriction of PCA vessels by 5-HT, we examined the effect of DAPK3 inhibition on intraluminal pressure-dependent myogenic reactivity. Vessels were isolated and subjected to three consecutive series of pressure steps (10 – 120 mmHg): first in normal Krebs buffer, then in Krebs solution containing 10 μM HS38 or DMSO vehicle control, and finally in zero external calcium (0 Ca2+) Krebs buffer. In control vessels, myogenic constrictions were initiated at 20 – 40 mmHg (**Figure 2A, panel i**), with maximal active constrictions at 120 mmHg of 148 μm or approximately 60% of the maximal vessel diameter under 0 Ca2+ conditions. These results agree with previously published myography studies of the PCA (Kim et al., 2013) and the MCA vessel (Johnson et al., 2009). DMSO had no effect on the myogenic response, neither the % dilation nor the active constriction was significantly impacted (**Figure 2A, panels ii and iii, respectively**). HS38 treatment markedly reduced the magnitude of myogenic constriction (**Figure 2B, panel i**), over a broad range of both low and high intraluminal pressures. The HS38-treated vessels retained 75.4 ± 4.3% of the maximal diameter measured under 0 Ca2+ conditions (**Figure 2B, panel ii**), and the average change in active constriction was 72.7 μm at 100 mmHg when control and HS38-treated vessels were compared (**Figure 2B, panel iii**).

**Figure 2.**
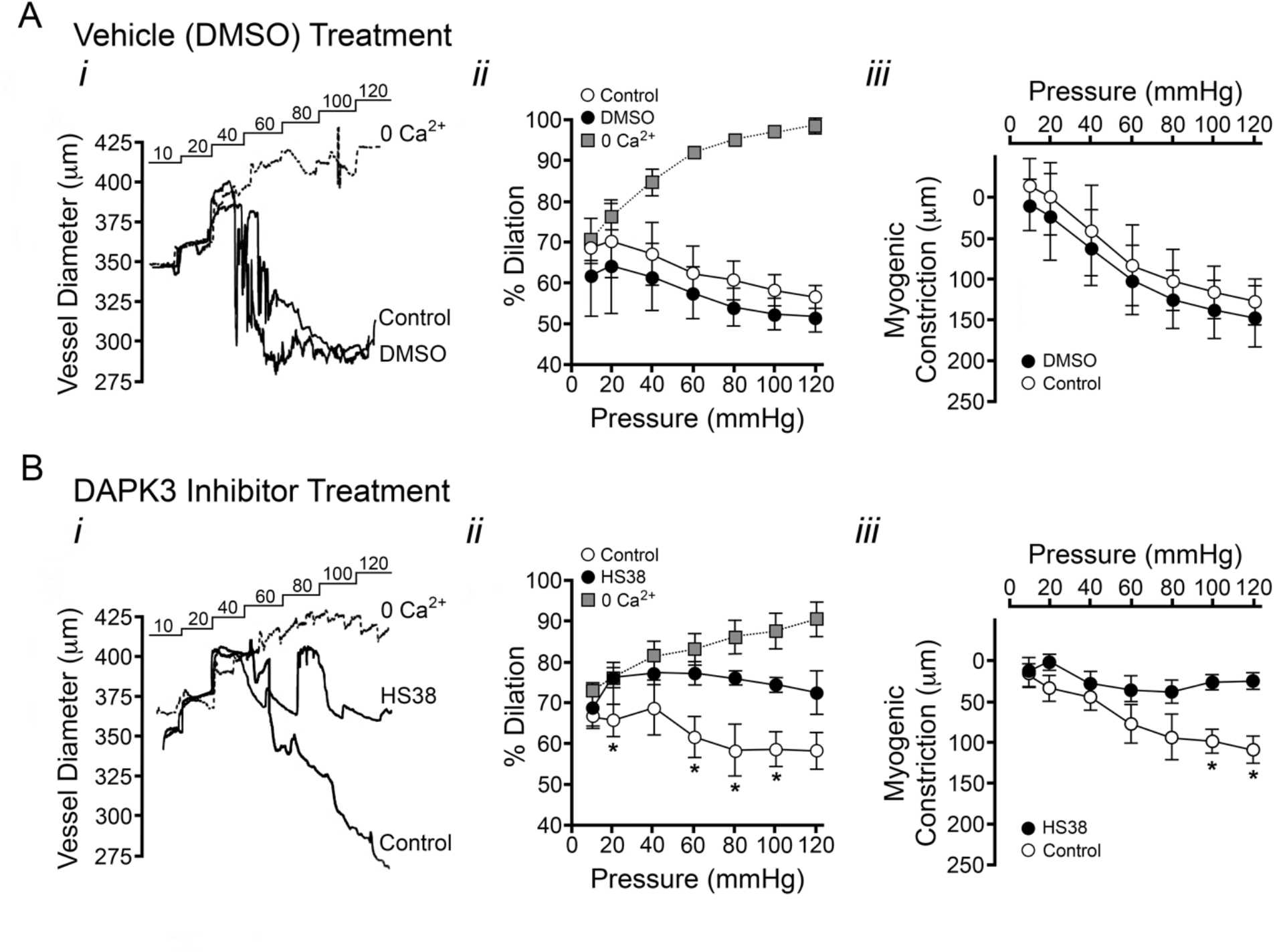
HS38 administration attenuates the myogenic response of rat posterior cerebral arteries. The myogenic reactivity of posterior cerebral arteries was monitored in the absence (**A**, DMSO vehicle) or presence of DAPK3 inhibitor (**B**, HS38, 10 μM). Data panels illustrate: (*i*) representative recordings of the change in outer diameter for vessels subjected to sequential pressure steps (10 – 120 mmHg) in the presence of normal Krebs buffer (NB), vehicle or DAPK3 inhibitor, and then calcium-free buffer (0 Ca2+); (*ii*), cumulative data showing vessel constriction relative to the maximum passive diameter observed in 0 Ca2+ solution (mean ± SEM, vessels obtained from n = 3 animals); and *(iii)* the magnitude of active myogenic constriction. *-significantly different from vessel constriction observed in the absence of treatment (two-way ANOVA with Sidak’s *post hoc* test, p < 0.05).

To determine if DAPK3 inhibition with HS38 has the potential to translate to the human vasculature, we assessed DAPK3 protein abundance as well as the myogenic response in set of human cerebral arteries obtained from surgical samples of four different patients. Importantly, immunoreactivity for DAPK3 was detected in all samples examined (**Figure 3A**), suggesting HS38 may impact vascular contractility in these vessels. Myogenic responses were recorded in vessels isolated from three of the four tissue biopsies (**Figure 3B**). Basal myogenic tone varied somewhat for the three vessels. This heterogeneity was expected as vessels were obtained from male and female patients of different ages with varied indications for surgery (e.g., epilepsy focal resection or tumor resection). The analysis of cumulative data showed the maximal myogenic constriction of human pial vessels to be 71.1 ± 6.8% of the passive vessel diameter (**Figure 3C**). Myogenic tone was significantly inhibited with application of HS38, reducing maximal constriction to 94.9 ± 3.9% of maximal vessel diameter at 120 mmHg and active myogenic responses by approximately 50 μm (**Figure 3D**).

**Figure 3.**
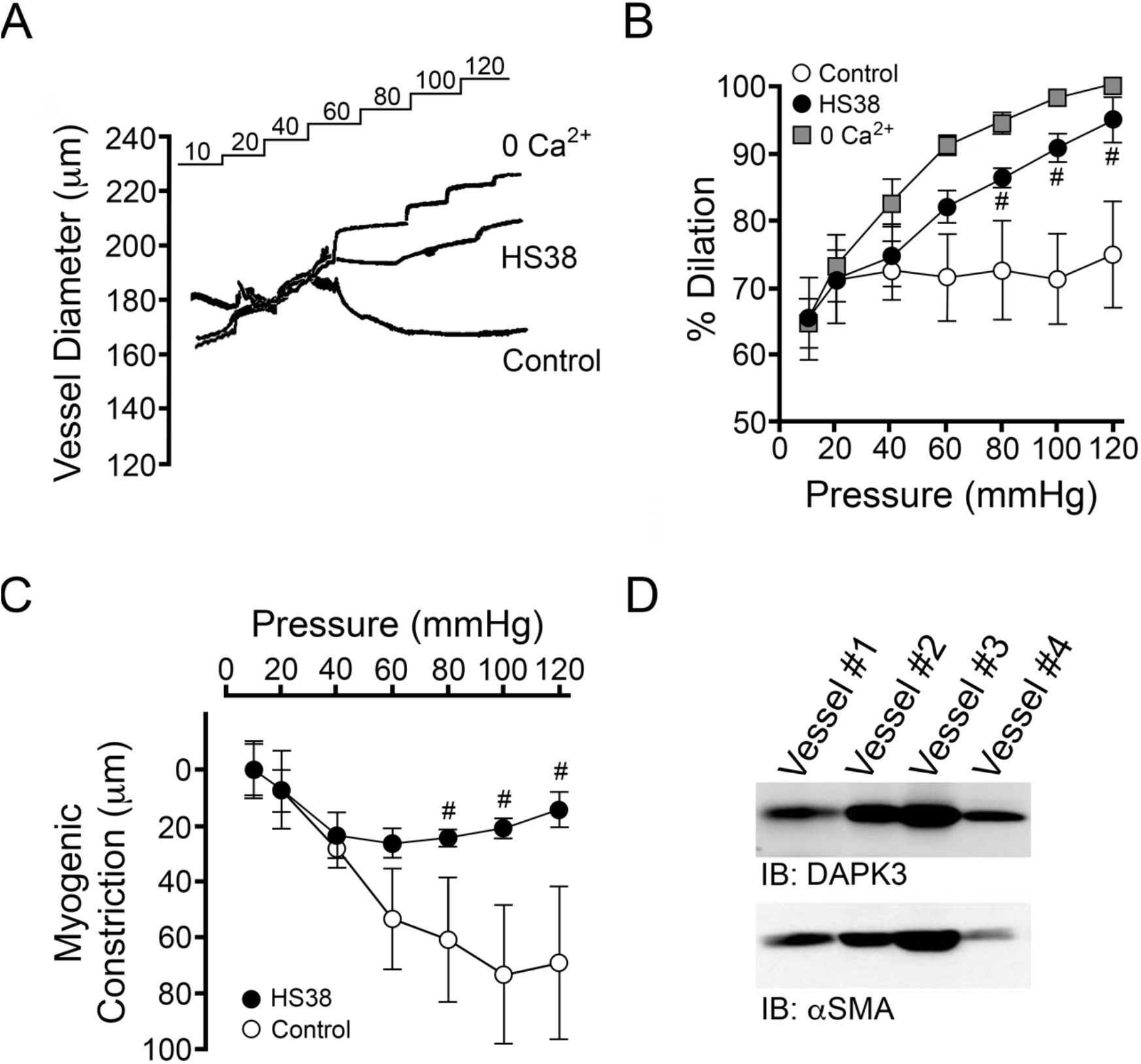
Myogenic responses of human cerebral arterioles are attenuated with HS38 treatment. **(A)**, representative recordings of human pial arterioles, outer vessel diameters, subjected to sequential 10 – 120 mmHg pressure steps in normal Krebs buffer (Control), with DAPK3 inhibitor (HS38, 10 μM), and in calcium-free Krebs buffer (0 Ca2+). Cumulative data are provided to show the vessel constriction relative to the maximum passive diameter observed in 0 Ca2+ solution **(B)** and the magnitude of active myogenic constriction **(C)**. Data are presented as mean ± SEM for n = 3 different experiments. Results were analyzed with two-way ANOVA and Sidak’s multiple comparisons test; #-significantly different from control vessel constriction observed in the absence of HS38, p < 0.05. **(D)**, pial arteries (~300-500 μm in diameter) were collected from four unique human brain tissue samples and immunoblotted for DAPK3 and smooth muscle actin (aSMA) as a loading control.

It was noted that cerebral vessels treated with HS38 did not appear to dilate fully and continued to exhibit some constriction during the final round of pressure steps in 0 Ca2+ Krebs buffer, relative to control vessels. Subsequently, rather than standardize data to the maximum passive diameter in 0 Ca2+ Krebs buffer at the end of the experiment as is exemplar in the literature (Schjorring et al., 2015; Wenceslau et al., 2021), a brief pressure step to 120 mmHg was performed immediately after mounting the vessel, and maximum vessel diameter was recorded prior to the development of any myogenic constrictions. To further interrogate the observed difference in passive vessel dilation following exposure to HS38, the intraluminal pressure-dependent myogenic responses of HS38-treated and DMSO vehicle control vessels were compared in the presence of 0 Ca2+ buffer (**Figure 4A**). HS38-treated vessels displayed distinct passive diameter responses to increased intraluminal pressure (p= 0.0002, two-way ANOVA), showing lower maximal diameters at pressures at/above 60 mmHg (**Figure 4B**).

**Figure 4.**
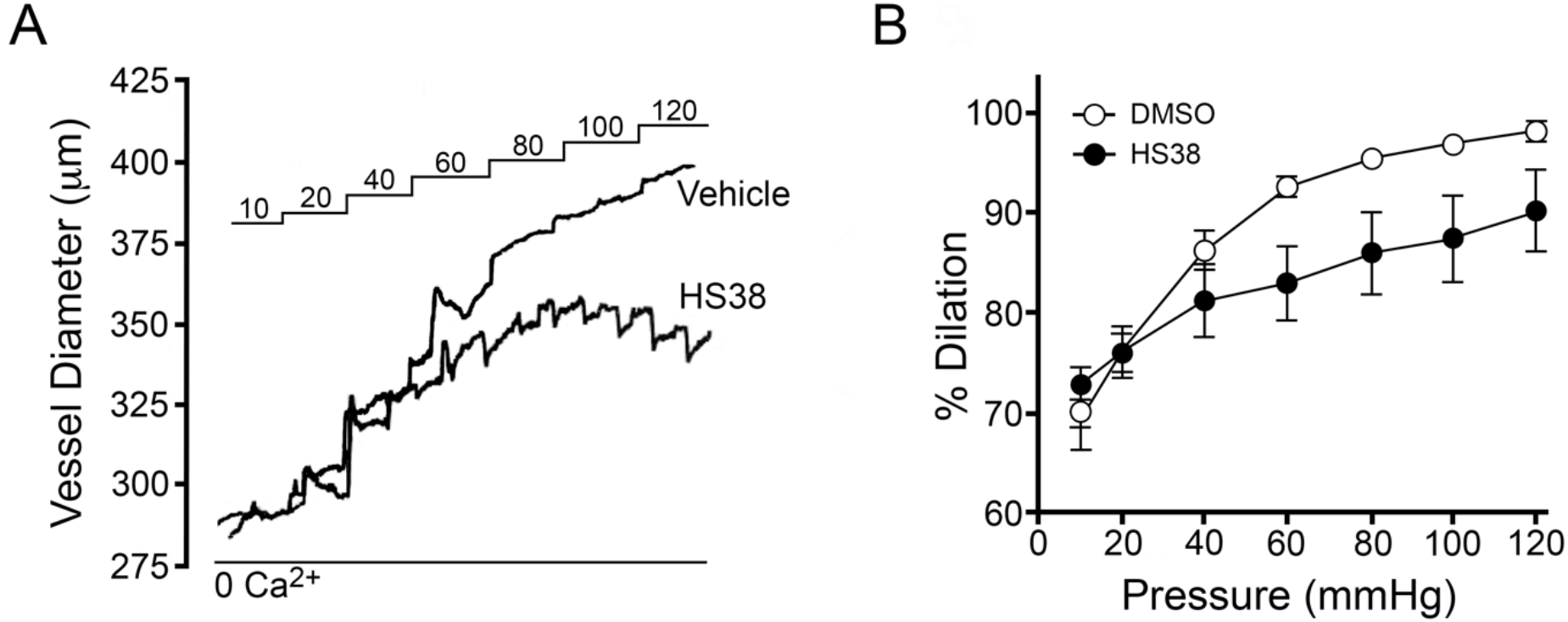
HS38 treatment of rat posterior cerebral arteries elicits increased passive diameters. **(A)**, representative recordings of the change in outer diameter for vessels subjected to sequential pressure steps (10 – 120 mmHg) in calcium-free Krebs buffer (0 Ca2+) following treatment with vehicle (DMSO) or DAPK3 inhibitor (HS38, 10 μM). Arteries were incubated at 10 mmHg for 30 min in the presence of DMSO or HS38, and then pressure steps (20 – 120 mmHg) were developed over an additional 60 min. In **(B)**, cumulative data show the vessel constriction relative to the maximum passive diameter observed in 0 Ca2+ solution (mean ± SEM, vessels obtained from n = 4 different animals). Groups were found to be significantly different (two-way ANOVA, p = 0.0002).

As seen in **Figure 5A**, the dilation afforded by DAPK3 inhibition with HS38 was not sustained indefinitely. When PCA vessels were held at an intraluminal pressure of 80 mmHg, the pharmacologic action of HS38 was characterized by a biphasic response: first, a rapid inhibition of myogenic tone and nearly complete dilation of the pressurized vessel within 5 minutes; and second, a slower recovery of myogenic tone and restoration of the original vessel diameter over the next 20 minutes (**Figures 5A, 5B**). As noted previously, some narrowing of vessels was also observed with HS38 in 0 Ca2+ solution when measuring the passive diameter at the experiment conclusion. Surprisingly, the dilation induced by HS38 treatment was not associated with any decline in the level of LC20 phosphorylation (**Figure 5C**). In contrast, the recovery of myogenic tone and vessel constriction was coupled with a significant increase in LC20 phosphorylation over that quantified either in the myogenic vessel prior to HS38 administration or in the maximally dilated vessel exposed to HS38. These results suggest that two distinct signaling pathways operate in the separate phases of the vessel response to HS38.

**Figure 5.**
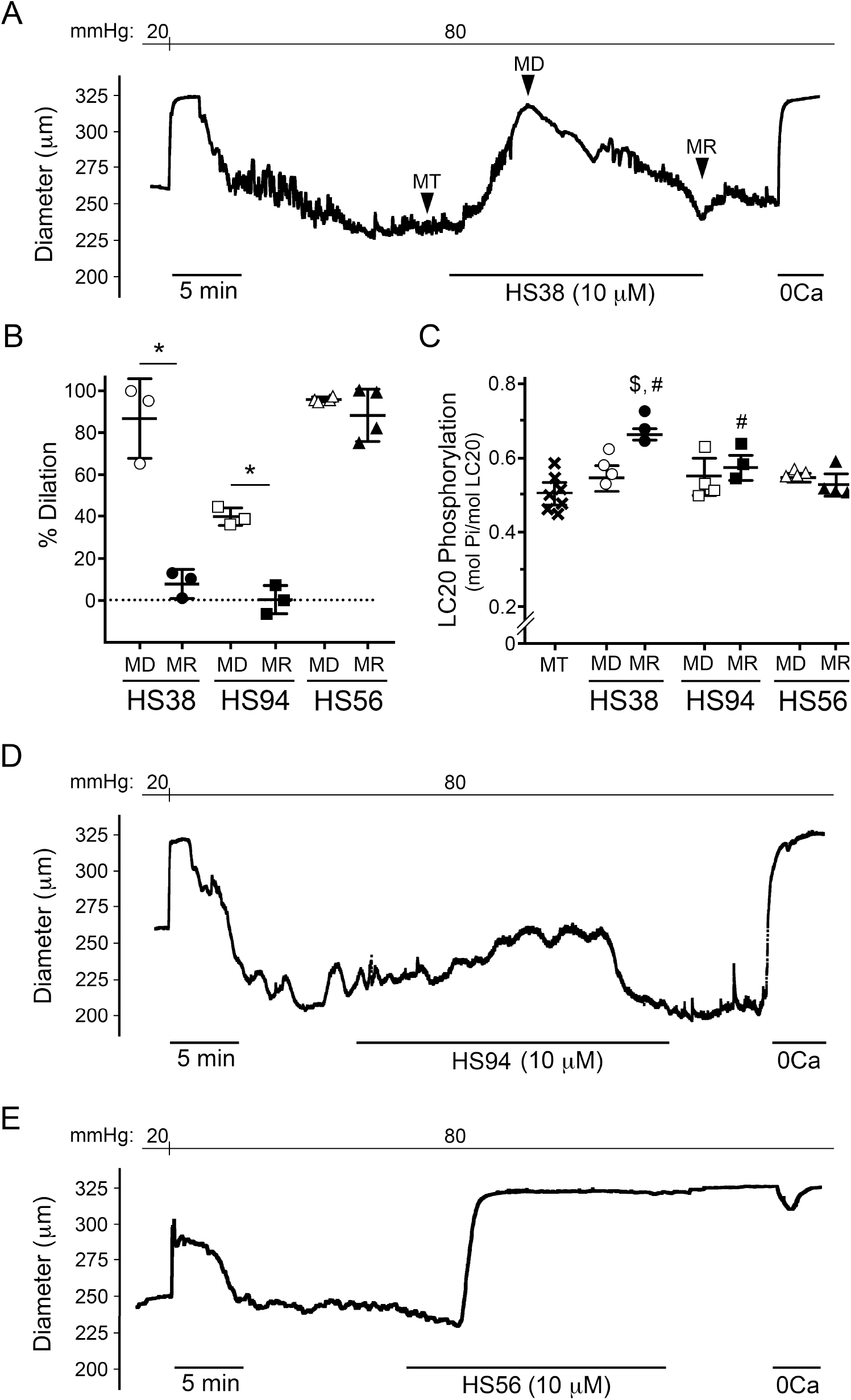
Prolonged HS38 treatment provides transient vasodilation of rat posterior cerebral arteries. **(A)**, representative vessel responses (outer diameter) to prolonged exposure with HS38 are shown. Vessels display pressure-dependent myogenic tone (MT) as well as vessel dilation (MD) and recovery of myogenic reactivity (MR) subsequent to HS38 exposure. In **(B)**, cumulative data show vessel dilation in response to HS38, HS94 and HS56 compounds; the % dilation at MD and MR were calculated relative to the myogenic response observed at MT and the passive diameter observed in 0 Ca2+ solution. Results were analyzed by Student’s t-test; *-significantly different from the vessel dilation (MD) observed with inhibitor treatment, p < 0.05. In **(C)**, the LC20 phosphorylation of vessels treated with HS38, HS94 and HS56 was quantified by Phos-tag SDS-PAGE. Results were analyzed by two-way ANOVA and Sidak’s multiple comparisons test; #-significantly different from vessels displaying myogenic tone (MT), p < 0.05; $-significantly different from the corresponding vessel dilated with the same inhibitor compound (MD), p<0.05. Representative recordings are also provided for the myogenic reactivity of vessels during exposure to HS94 **(D)** and HS56 **(E)**. Data are mean ± SEM; PCA vessels were obtained from n = 3-7 different animals.

HS38 has excellent potency and specificity for DAPK3 and, unlike other DAPK3 inhibitor compounds, exhibits no off-target actions on Rho-associated protein kinase (ROCK) (Al-Ghabkari et al., 2016). Although HS38 is highly specific for DAPK3 when compared to other contractile kinases in smooth muscle (Carlson et al., 2013), the compound does possess equal efficacy for the Provirus integrating site Moloney murine leukemia virus (PIM3) kinase. To differentiate the DAPK3-mediated phenotype from the PIM3-mediated phenotype, additional myography was conducted with second generation compounds based on the same molecular scaffold (Carlson et al., 2018): HS94 (less inhibitory activity towards PIM3) and HS56 (more inhibitory activity towards PIM3). As shown in **Figure 5D**, HS94 yielded a similar phenotype as HS38 in that a biphasic response was observed with vessel dilation and then recovery of myogenic tone over time (**Figure 5B**). Again, the dilation response to HS94 administration was not coupled to changes in LC20 phosphorylation whereas an increase in LC20 phosphorylation was found with the recovery of myogenic tone (**Figure 5C**). However, the kinetics of dilation and recovery were distinct from those observed with HS38 with a slower rate of vessel dilation and a more rapid recovery of vessel constriction. Finally, HS56 treatment elicited a novel phenotype (**Figure 5E**), wherein rapid and complete dilation of the PSA vessels occurred without recovery of myogenic tone (**Figure 5B**) or alteration in LC20 phosphorylation status (**Figure 5C**). In this case, the dilatory response of vessels to HS56 was irreversible.

## DISCUSSION

This is the first study to identify a role for DAPK3 in the myogenic response of the cerebral resistance vasculature. We utilized the HS38 compound, a specific DAPK3 inhibitor that does not target other VSM contraction-related kinases (Carlson et al., 2013; Al-Ghabkari et al., 2016; Carlson et al., 2018), to demonstrate the contribution of DAPK3 to the myogenic reactivity of PCA vessels. Additional evidence supports the translational importance of DAPK3 to the human cerebral vasculature, with robust expression of the protein kinase and significant HS38-dependent attenuation of myogenic reactivity observed for pial vessels. Finally, intriguing pharmacokinetics are observed for a suite of DAPK3 inhibitors that suggest intrinsic compensation of additional signaling pathways in response to attenuation of DAPK3 activity as well as a putative novel impact of PIM3 in the myogenic response of cerebral arterioles.

The vascular myogenic response is a complex and highly regulated process that is the focus of much investigation (Hong et al., 2020; Jackson, 2021). Myogenic constrictions, which can occur in isolated endothelium-denuded vessels in response to increasing luminal pressure, have a Ca2+ sensitive component, a Ca2+ sensitization component and an actin cytoskeletal reorganization component (Osol et al., 2002; Cole and Welsh, 2011; Walsh and Cole, 2013; El-Yazbi and Abd-Elrahman, 2017). Considerable evidence links DAPK3 to mechanisms of Ca2+ sensitization in smooth muscle contraction (Haystead, 2005; Ihara and MacDonald, 2007); namely, through the Ca2+-independent phosphorylation of myosin regulatory light chain (LC20) or by the inhibition of myosin phosphatase (i.e., either by direct phosphorylation of the myosin phosphatase targeting subunit MYPT1 or by the phosphorylation of the myosin phosphatase inhibitor CPI-17). Intriguingly, the dilatory responses of PCAs to HS38 occurred in the absence of corresponding changes to LC20 phosphorylation. The lack of change in LC20 phosphorylation with HS38 excludes contractile mechanisms involving 1) a change in Ca2+ influx and cytosolic free [Ca2+] due to altered ion channel activity (e.g., decreased Ca2+ channel opening or increased K+ channel opening and hyperpolarization leading to indirect decline in cytosolic [Ca2+] and contraction; indeed, HS38 was previously identified to have no effect on the depolarization-induced Ca2+ transient (MacDonald et al., 2016)), or 2) a change in myosin phosphatase activity through the inhibitory actions of ROCK or CPI-1, as both would result in a decrease in LC20 phosphorylation. This, as well as the observation that HS38 exposure elicits narrowing of PCA diameters under 0 Ca2+ conditions and high intraluminal pressures, suggests that DAPK3 does not regulate Ca2+ sensitization pathways during the myogenic response of PCA vessels but rather operates mechanistically to control the actin cytoskeleton.

There is acknowledgement for a critical role of actin cytoskeleton remodeling in the development of myogenic tone (Cole and Welsh, 2011; Walsh and Cole, 2013; Colinas et al., 2015; Hong et al., 2016). Indeed, mechanisms that provide Ca2+ sensitization of LC20 phosphorylation and force development may not be the only means for DAPK3 contributions to the myogenic character of PCA vessels. DAPK3 is known to contribute to actin polymerization and focal adhesion dynamics with impact on the motility of vascular smooth muscle cells and the contraction of isolated smooth muscle cells (Komatsu and Ikebe, 2004). DAPK3 overexpression in fibroblast cells was associated with actin cytoskeletal remodeling, expansion of focal adhesions and decreased focal adhesion kinase (FAK) Tyr397 phosphorylation (Nehru et al., 2013), and increased FAK Thr297 phosphorylation was required for the pressure-dependent myogenic responses of cerebral arterioles (Colinas et al., 2015). However, comprehensive examinations of the role played by DAPK3 in the actin cytoskeletal dynamics during contraction of isolated VSM or during the myogenic responses of resistance vessels remains to be completed. For example, additional insight is required to understand the mechanism for the slow return of tone associated with greater LC20 phosphorylation during the sustained exposure of cerebral arterioles to DAPK3 inhibitors.

RhoA/ROCK signaling is also involved in the actin polymerization that accompanies myogenic constrictions of cremaster arterioles (Moreno-Dominguez et al., 2013), which may indicate a role for DAPK3 downstream of ROCK. In non-smooth muscle cells, ROCK can phosphorylate DAPK3 at Thr265 and Thr299, leading to its activation and cytosolic localization (Hagerty et al., 2007). Considering the potent inhibitory effect of ROCK-specific inhibitors on the myogenic response (Johnson et al., 2009; Moreno-Dominguez et al., 2013), it is possible that DAPK3 may be involved downstream of ROCK to influence the remodeling of the actin cytoskeleton during myogenic constriction, and this relationship will require further investigation. In addition, DAPK3 was positioned downstream of G protein-coupled receptors (GPCRs) that signal through Gq/11 and/or G12/13, and findings implied that the activation of RhoA/ROCK was required for subsequent DAPK3-dependent events (MacDonald et al., 2016). These signaling linkages coincide with emerging evidence that identify the angiotensin II receptor (AT1R) and Gq/11-coupled signals to function as sensors of membrane stretch in VSM cells (Mederos y Schnitzler et al., 2008; Kauffenstein et al., 2012; Hong et al., 2016; Pires et al., 2017). With specific DAPK3 and ROCK inhibitors now available, the mechanoactivation of AT1R and subsequent signaling through ROCK and DAPK3 should be thoroughly evaluated in relation to the myogenic response.

The HS56 compound was previously used to characterize polypharmacology of DAPK3 and PIM3 inhibition in vascular smooth muscle systems (Carlson et al., 2018). Indeed, acute decreases in systolic blood pressure (SBP), likely elicited as a coincident impact of peripheral vasodilation, were observed following intravenous administration of HS56 into wild-type and spontaneously hypertensive renin transgene (RenTg) mice. In the RenTG mice, HS56 infusion elicited dose-dependent decreases in SBP for the duration of monitoring (~2.5 min) with a drop in SBP of ~30 mmHg. While HS38 and HS94 were not assessed in mice, another analogue (i.e., HS148; with high degree of selectivity for DAPK3 over PIM3) did not affect SBP upon infusion. We expect that compensatory signaling mechanisms may have masked the dilatory impact of *in vivo* DAPK3 inhibition. The lack of physiological impact of HS148 on SBP in mice correlates with the transient nature of dilation observed with selective inhibition of DAPK3 *ex vivo* in PCAs and suggests that polypharmacology of PIM3/DAPK3 acts in a fundamentally different manner (Sawada et al., 2006).

Our findings also establish PIM3 as a novel signaling molecule in the myogenic process that is worthy of further investigation. The PIMs are considered oncogenic Ser/Thr protein kinases with an expansive scope of biological impact on cell growth, proliferation and survival, cell differentiation, apoptosis and metabolism (Sawada et al., 2006). More recent studies demonstrate PIM involvement in cancer cell migration and metastatic invasion. In endothelial cells, PIM3 silencing attenuated cellular migration and vascular tube formation during angiogenesis; PIM3 was also shown to colocalize with FAK at the lamellipodia (Zhang et al., 2009). In excised rat caudal artery, HS56 treatments were found to suppress Ca2+ sensitization and reduce smooth muscle force generation with attenuation of LC20 phosphorylation. *In vitro* biochemical assessments identified MYPT1 to be an effective substrate target of PIM3 although the specific phosphorylation sites remain to be mapped. Ultimately, it remains to be determined whether PIM3 also impacts upon the actin cytoskeleton.

**Table 1.** Inhibitor constants for HS compounds. Data extracted from Carlson *et al.* (Carlson et al., 2018) The relative inhibitory potency was judged to be: DAPK3 = PIM3 for HS38 (1:1), DAPK3>>>PIM3 for HS94 (20:1) and DAPK3 < PIM3 for HS56 (1:4).

| | $K_i$ (μM) | | |
| --- | --- | --- | --- |
|  | HS38 | HS94 | HS56 |
| DAPK3 | 0.260 | 0.126 | 0.315 |
| PIM3 | 0.208 | 2.50 | 0.072 |

## Acknowledgements

The authors thank the members of the Calgary Stroke Program who enabled the collection of tissue samples from patient donors.

## Author Contributions

S.R.T., M.C., and C.S. completed the data analysis and prepared figures. D.A.C synthesized the HS38, HS94 and HS56 compounds. T.A.J.H. coordinated the production of inhibitor compounds and made intellectual contributions to the project. W.C.C. made intellectual contributions to the project. J.A.M. conceived and coordinated the study, assisted with experimental design, wrote the manuscript, provided trainee supervision, and made intellectual contributions to the project. All authors reviewed the results and approved the final version of the manuscript.

